# Widespread gray and white matter microstructural alterations in dual cognitive-motor impairment

**DOI:** 10.1101/2025.04.18.649603

**Authors:** Kavita Singh, Yang An, Kurt G. Schilling, Dan Benjamini

**Affiliations:** Multiscale Imaging and Integrative Biophysics Unit, National Institute on Aging, NIH, 251 Bayview Blvd., Baltimore, MD 21224, USA; Brain Aging and Behavior Section, National Institute on Aging, NIH, 251 Bayview Blvd., Baltimore, MD 21224, USA; Department of Radiology and Radiological Sciences, Vanderbilt University Medical Center, 1211 Medical Center Drive, Nashville, TN 37232

**Keywords:** **Keyword:** Dual cognitive-motor impairment, HCP-Aging, temporal-meta ROIs, Mean apparent propagator MRI, gray matter microstructural, white matter motor tracts

## Abstract

**INTRODUCTION:** Dual cognitive-motor impairment in aging is a strong predictor of dementia, yet its effects on vulnerable gray matter regions microstructure remain unexplored.

**METHODS:** This study classified 582 individuals aged 36–90 into cognitive-motor impairment, isolated cognitive or motor impairment, and control groups. Microstructural differences in 27 temporal and motor-related gray matter regions and white matter tracts were assessed using DTI and mean apparent propagator (MAP-MRI), a technique well-suited for gray matter analysis.

**RESULTS:** We found widespread microstructural alterations in gray and white matter among individuals with dual cognitive-motor impairment. These changes were not observed in isolated cognitive or motor impairment after multiple comparisons correction.

**DISCUSSION:** Dual cognitive-motor impairment is associated with reduced cellular density in temporal gray matter, decreased fiber coherence, and potential demyelination in white matter tracts, suggesting widespread microstructural disruption. These findings could help understand brain aging and facilitate interventions to slow neurodegeneration and delay dementia onset.

## 1. Background

Cognitive function declines with aging, with approximately 40% of individuals over 65 years experiencing some form of memory loss [1]. While aging is a well-established risk factor for neurodegenerative diseases such as Alzheimer’s disease (AD) and dementia, prodromal stages— such as mild cognitive impairment (MCI) and motoric cognitive risk syndrome (MCR)—are often overlooked, with only 11.4% of individuals receiving a timely diagnosis for the latter [2]. In some cases, motor impairment precedes cognitive decline and serves as a strong predictor of dementia, affecting approximately 9.7% of older adults [3]. Evidence across diverse populations suggests that individuals exhibiting dual decline in both cognition and mobility face a markedly higher risk of developing dementia than those with deficits in either domain alone [4–6]. This co-occurrence—characterized by simultaneous declines in gait speed and memory—may represent a distinct clinical phenotype with strong prognostic value for identifying individuals at elevated risk for dementia. It is not yet known whether synergistic impairment in cognition and mobility reflects brain biomarker patterns similar to those seen in early AD and related neurodegenerative conditions. Detecting biomarker changes linked to dual decline or impairment may offer key insights into its underlying mechanisms and its potential role in dementia development.

Recent MRI studies have investigated the underlying cerebral changes in distinct patterns of cognitive-motor impairment, however, these studies have primarily focused on macrostructural or volumetric alterations, such as hippocampal atrophy, reductions in total gray matter (GM) volume, and ventricular enlargement [4]. Microstructural changes in white matter (WM) have been investigated; however, these studies primarily assessed global alterations, such as WM hyperintensities and total WM volume [7], lacking regional specificity. Notably, tau PET-derived temporal meta regions of interest (ROIs) are among the earliest affected in clinically normal individuals with advancing age [8] and also in preclinical AD population [9] , and should therefore be analyzed alongside motor-related regions to fully capture subtle microstructural changes due to cognitive-motor impairment.

Microstructural alterations in GM are primarily investigated using ex vivo histological analysis, as current MRI techniques have limited sensitivity and modeling capabilities for detecting in vivo GM changes[10,11]. While in vivo MRI studies have successfully identified WM alterations using diffusion tensor imaging (DTI) [12], this approach remains insufficiently sensitive for assessing microstructural changes in GM [13]. The emergence of probabilistic models such as the mean apparent propagator (MAP) [14] —a diffusion MRI (dMRI) framework that operates independently of biophysical assumptions—has enabled the use of multi-shell dMRI data to investigate gray matter microstructural changes associated with the AD pathological cascade [15] and aging[16]. MAP-MRI was recently utilized to analyze a large cross-sectional dataset from the Human Connectome Project-Aging (HCP-A), which included 725 subjects aged 36 to 90 years. The findings revealed that MAP-MRI features strongly correlated with memory and executive functions across various cortical and subcortical GM regions [13]. However, the HCP-A cohort exhibited concerning Montreal Cognitive Assessment (MoCA) [17] scores of 24 or below in nearly 15% of subjects.

Building on this evidence, we hypothesize that cognitive-motor impairment affects gray and white matter cerebral microstructure and that these changes can be detected using MAP-MRI and DTI metrics. This study aims to (1) classify HCP-Aging subjects into groups based on probable cognitive impairment (CI), motor impairment (MI), cognitive-motor impairment (CMI), and unimpaired or cognitive-motor normal (CMN); (2) validate these findings by assessing group differences in established MoCA scores and walking speed measures; and (3) investigate microstructural alterations in gray and white matter by analyzing MAP-MRI and DTI metrics in temporal meta-ROIs, motor-related GM regions, and associated WM tracts. Through this approach, we seek to foreword underlying microstructural changes in cognitive-motor impairment in aging population.

## 2. Methods

### 2.1 Study data and participants

Data were obtained from the Human Connectome Project-Aging (HCP-A) Lifespan 2.0 Release, comprising 725 healthy adults aged 36 to 90 years, collected across four sites using harmonized MRI acquisition protocols [14]. The study was approved and overseen by the Institutional Review Board, and written informed consent was obtained from all participants. While no individuals were excluded based on medication use, self-reported medication data were collected during study visits to identify or account for potential confounding effects.

Data were obtained from the Human Connectome Project-Aging (HCP-A) Lifespan 2.0 Release, comprising 725 healthy adults aged 36 to 90 years, collected across four sites using harmonized MRI acquisition protocols [18]. The study was approved and overseen by the Institutional Review Board, and written informed consent was obtained from all participants. While no individuals were excluded based on medication use, self-reported medication data were collected during study visits to identify or account for potential confounding effects. Participants with a history of major psychiatric disorders (e.g., schizophrenia, bipolar disorder) or neurological conditions (e.g., stroke, brain tumors, Parkinson’s disease) requiring diagnosis and treatment were excluded based on essential health assessments Of 725 healthy subjects’ data available, we included 582 subjects for MAP-MRI and DTI analysis based on quality control of the raw data. The study population demographics is provided in Table 1. During HCP-A 2.0 release, data from participants over the age of 90 years were grouped into a single age bin, precluding precise age-specific analyses due to data policy, and therefore were excluded from our study. Years of education, body mass index were also recorded and used in the final analysis.

**Table 1.** Participant demographics (N = 582) and list of neurobehavioral test scores expressed as mean (standard deviation).

| **Groups** | **Cognitive motor normal (CMN)** | **Cognitive impairment (CI)** | **Motor impairment (MI)** | **Cognitive motor impairment (CMI)** | **Post hoc statistical analysis** |
| --- | --- | --- | --- | --- | --- |
| Age, mean±SD, [years] | 58.00±14.3 | 59.88±14.8 | 57.74±14.03 | 58.24±13.3 | #1 |
| Males/Females | 65/48 | 88/52 | 57/138 | 42/92 | #2 |
| Education [years] | 18.19±2.5 | 17.52±1.9 | 17.77±2.0 | 16.51±1.9 | #3 |
| NIH Fluid | 109.95±9.8 | 92.86±8.9 | 105.66±8.7 | 88.87±8.4 | #4 |
| Walk Endurance | 708.09±65.2 | 661.26±65.2 | 578.70±65.2 | 532.85±80.6 | #5 |
| MoCA | 27.73±2.0 | 25.64±2.5 | 27.16±2.1 | 25.22±2.5 | #6 |
| Walk Speed [m/s] | 1.35±0.2 | 1.27±0.2 | 1.27±0.2 | 1.18±0.2 | #7 |

#1 - ANOVA showed no significant difference among groups
#2 - Chi-square result showed significant differences in CMN vs CI, CMN vs CMI, MI vs CI, and MI vs CMI
#3 - ANOVA showed significant differences in CMN vs CI, CMN vs CMI, CMI vs MI, and MI vs CMI
#4 - ANOVA showed significant differences among all groups.
#5 - ANOVA showed significant differences among all groups.

### 2.2 Participant classification and its validation

The HCP-A conducted detailed behavioral and cognitive assessments using NIH toolbox [19]. The tests and respective scores that were used in the current study are provided in Table 1. Cognitive and physical function were assessed using two standardized NIH Toolbox measures. First, the NIH Fluid Cognition Composite Score was derived from five component tasks—Flanker Inhibitory Control and Attention, Dimensional Change Card Sort, Picture Sequence Memory, List Sorting Working Memory, and Pattern Comparison Processing Speed. This composite score evaluates key domains such as problem-solving, cognitive flexibility, processing speed, and episodic memory formation, which are particularly sensitive to age-related neurobiological changes and are commonly affected in neurodegenerative disorders. Second, cardiovascular endurance was measured using the NIH Toolbox 2-Minute Walk Test, in which participants walked a 10-foot out-and-back course for two minutes, and total distance was recorded. This measure captures gradients in physical performance, even among older adults with unimpaired gait speed or no self-reported mobility limitations [20]. Participants were categorized into four groups based on their performance in both tests using k-means clustering (as detailed in the Statistical Analysis). The groups included cognitive-motor normal (CMN), cognitive impairment (CI), motor impairment (MI), and cognitive-motor impairment (CMI).

The Montreal Cognitive Assessment (MoCA) has been widely used to detect cognitive impairment, including mild cognitive impairment (MCI) and early Alzheimer’s disease [17], as well as in individuals with Parkinson’s disease [21]. Similarly, reduced walking speed— commonly observed with aging—has emerged as a robust predictor of adverse health outcomes, including increased risk of falls and mortality [22]. These two measures have been used to classify individuals exhibiting concurrent cognitive and motor decline, referred to as dual decline [23]. In the present study, we use MoCA composite scores and gait speed to validate the categorization of participants into cognitive, motor, or dual impairment groups.

### 2.3 Imaging data acquisition

To achieve the recruitment and diversity objectives of the Human Connectome Project in Aging (HCP-A), imaging data were acquired across four participating institutions using harmonized protocols. All sites employed a 3T Siemens MAGNETOM Prisma scanner equipped with a 32- channel head coil.

Structural MRI included both T1-weighted (T1w) and T2-weighted (T2w) imaging. T1w images were acquired using a multi-echo MPRAGE sequence with 0.8 mm isotropic resolution (TR = 2500 ms, TI = 1000 ms, TE = 1.8/3.6/5.4/7.2 ms, flip angle = 8°). T2w images were obtained using a T2w-SPACE sequence with 0.8 mm isotropic resolution (TR = 3200 ms, TE = 56.4 ms). Both sequences incorporated embedded volumetric navigators for prospective motion correction and selective reacquisition of motion-corrupted k-space lines. For subsequent image processing, the average of the first two echoes from the MPRAGE acquisition was used.

Diffusion-weighted imaging (DWI) was conducted using a pulsed gradient spin-echo EPI sequence with 1.5 mm isotropic voxel size (TR/TE = 3230/89.5 ms). Diffusion encoding employed two b-value shells (1500 and 3000 s/mm²), each with 98–99 gradient directions, and 28 interleaved b=0 images. Data were acquired with both anterior–posterior (AP) and reversed (PA) phase encoding directions. Full acquisition details are provided in Harms et al. [18].

### 2.4 Data processing

#### 2.4a. Preprocessing

Each participant DWIs were manually quality checked before and during each processing step. The preprocessing modules used in this work are part of the TORTOISE dMRI processing package [24]. Briefly, the dMRI data initially underwent denoising with the MPPCA technique [25], which was followed by Gibbs ringing correction [26] for partial k-space acquisitions [27]. Motion and eddy currents distortions were subsequently corrected with TORTOISE’s DIFFPREP module [28] with a physically-based parsimonious quadratic transformation model and a normalized mutual information metric. For susceptibility distortion correction, a T2w image was fed into the DRBUDDI [24] method for phase-encoding distortion correction. The final preprocessed data was output with a single interpolation in the space of an anatomical image at native in-plane voxel size.

#### 2.4b. Gray matter and white matter brain regions of interest

The spatially localized atlas network tiles (SLANT) method was used to perform whole brain segmentation [29]. Briefly, SLANT employs multiple independent 3D convolutional networks for segmenting the brain resulting in the labels for 132 anatomical regions based on the BrainCOLOR protocol (https://mindboggle.info/braincolor/). SLANT is publicly available and has shown high intra- and inter-scan protocol reproducibility [30]. From the SLANT-derived 132 labels we included 13 regions, merged bilaterally, of which 7 were temporal-meta ROIs and 6 motor ROIs, as shown in Figure 1A. These labels were first transformed from T1w space to T2w/DWI space using Advanced Normalization Tool (ANTs, Philadelphia, PA, United States) rigid registration [31]. Additionally, all ROIs were eroded using a 2×2×2 voxels cubic structuring element to reduce partial volume effects and imperfect image registration and to mitigate structural atrophy seen especially at older ages.

**Figure 1:**
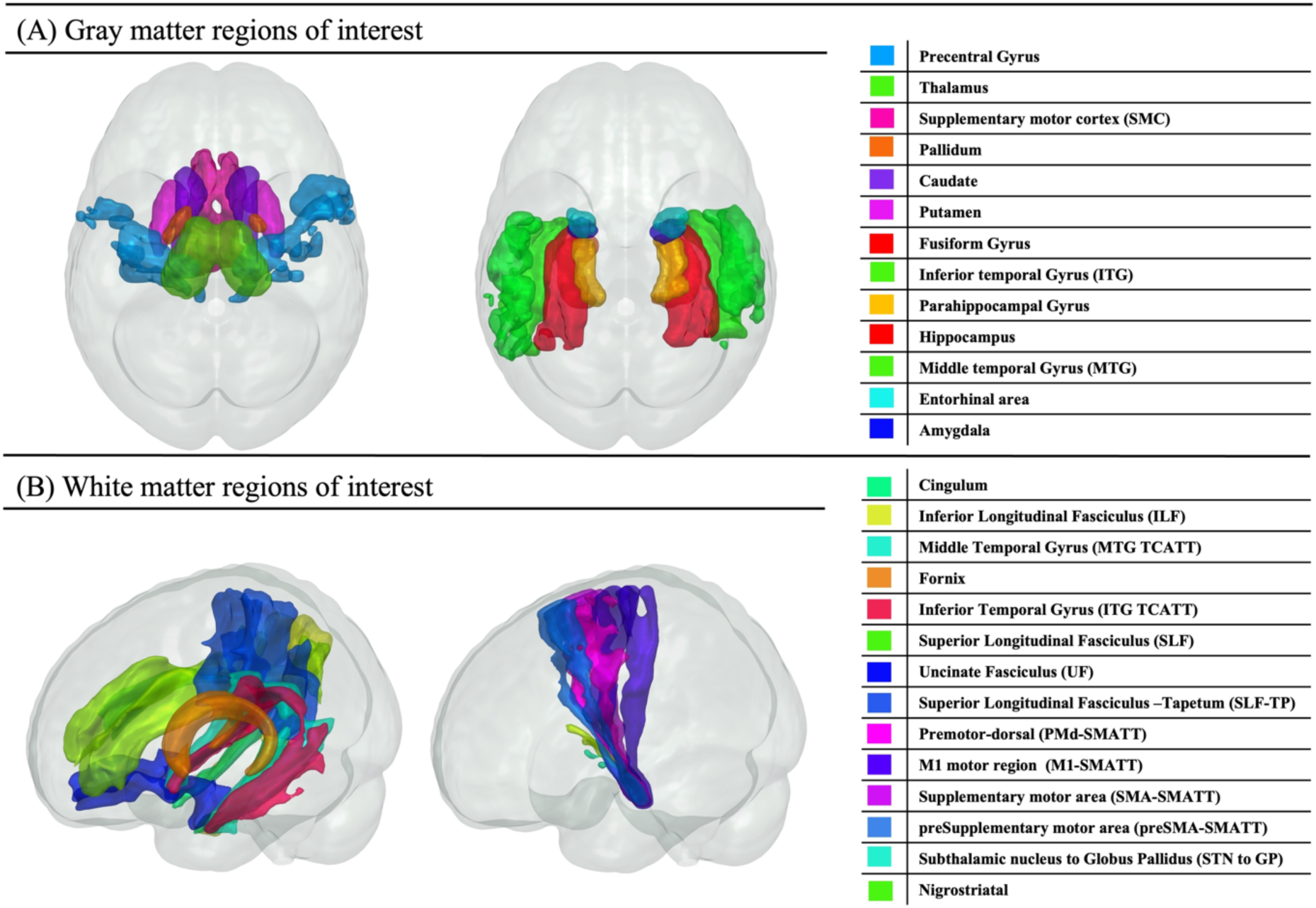
Investigated regions of interest. Three-dimensional rendering of (A) temporal meta-ROIs and motor-related GM regions, and (B) associated WM tracts. A total of 27 ROIs were investigated in the current study

To comprehensively investigate microstructural alterations in white matter (WM) pathways connected to gray matter (GM) regions of interest—specifically temporal meta-ROIs and motor-related areas—we analyzed 14 WM tracts identified as afferent or efferent projections to these regions [28]. Tract delineation was based on the Vanderbilt University Medical Center (VUMC) tractography atlas [29], which was developed using data from the HCP-Aging dataset and is illustrated in Figure 1

To comprehensively investigate microstructural alterations in WM pathways connected to GM regions of interest—specifically temporal meta-ROIs and motor-related areas—we analyzed 14 WM tracts identified as afferent or efferent projections to these regions [32]. Tract delineation was based on the Vanderbilt University Medical Center (VUMC) tractography atlas [33], which was developed using data from the HCP-Aging dataset and is illustrated in Figure 1B. Individual subject T1w images were nonlinearly registered using ANTS to a 2.0 mm isotropic MNI ICBM 152 asymmetric template [31]. Initial transformations of T2w/DWI space to T1w were concatenated with T1w to MNI transforms to bring subject data in native space to MNI 2mm space. Later, WM labels in MNI space were used to compute MAP-MRI and DTI average metrics in these four groups of subjects.

#### 2.4 c. MAP-MRI and DTI parameters estimation

Using the preprocessed DWIs, we estimated the voxel-wise diffusion propagators using a MAP-MRI series expansion truncated at order 4 [34]. MAP-MRI extends the DTI model, harnessing the capabilities of advanced dMRI sequences to offer potentially more sensitive metrics for detecting early pathological events in the disease process [35] and to quantify microscopic flow [36]. The commonly derived metrics are the propagator anisotropy (PA), which is a generalized version of DTI’s fractional anisotropy (FA); the non-Gaussianity (NG), which quantifies the dissimilarity between the propagator, and its Gaussian part; and the zero displacement probabilities, including the return to the origin probability (RTOP), the return to the axis probability (RTAP) and the return to the plane probability (RTPP), which comprehensively quantify various features of the three-dimensional diffusion process. The physical meaning and common biological interpretation of the MAP-MRI parameters are summarized in Table 2. Note that throughout the paper we report the RTAP^1/2^ and RTOP^1/3^ values to allow consistency in units of the zero-displacement probability metrics (i.e., 1/mm). The mean MAP metrics (NG, PA, RTAP, RTOP, and RTPP) from 27 temporal-meta and motor ROIs were computed. Using the same processing strategy, we computed fractional anisotropy (FA), axial diffusivity (AD), radial diffusivity (RD), trace (TR) using TORTOISE for 27 ROIs in each participant.

**Table 2.**
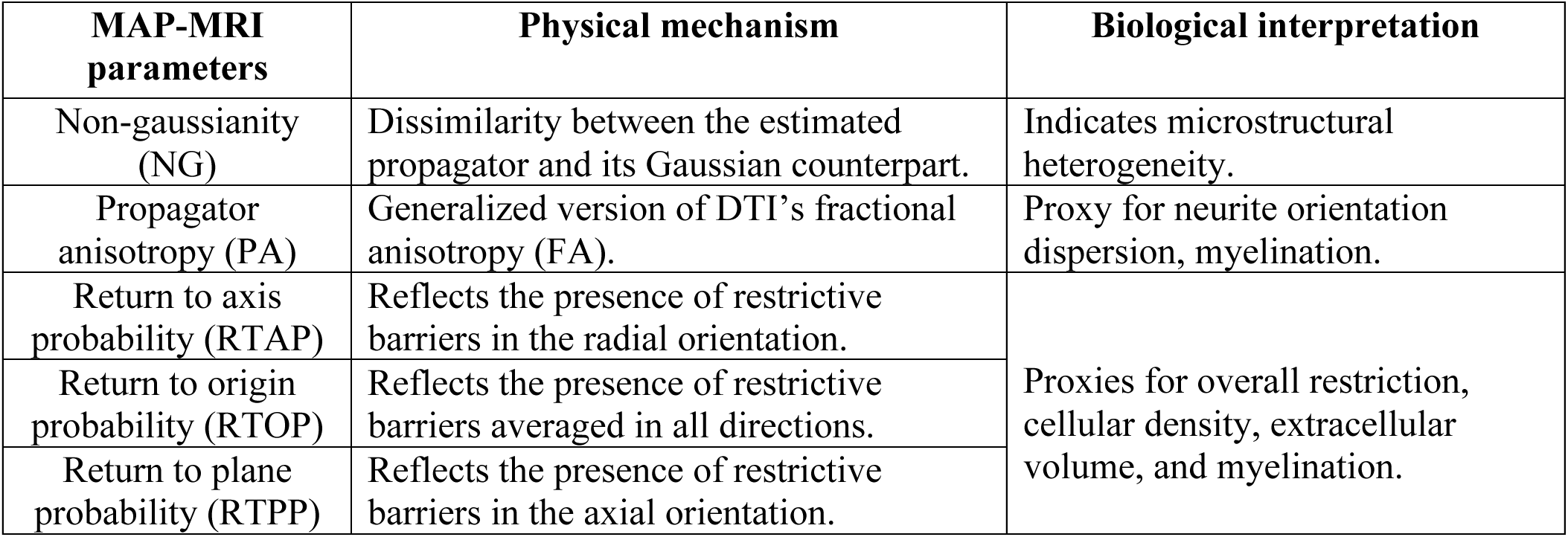
MAP-MRI parameters, their physical mechanism and common biological interpretations.

### 2.5 Statistical analysis

To account for age-related variability in cognitive and motor test scores, we performed separate multiple linear regressions with z-standardized test scores as the outcomes; age and age² as predictors; and extracted the residuals. These residuals represent age-adjusted relative scores, where positive values indicate performance above the age-related mean and negative values indicate performance below it. These adjusted scores were then used as inputs for k-means clustering to classify subjects into the four study groups. The Elbow Method was used to determine the optimal number of clusters (k=4) and initialize the algorithm. K-means clustering was then performed with a maximum of 1,000 iterations and 10 replicates. The resulting groups were analyzed for differences in age, and years of education using ANOVA (sex using Chi-square test), with Bonferroni correction applied for multiple comparisons.

To validate the grouping, we used MoCA and the 4 Meter Walk Speed scores as independent cognition and motor performance metrics. For MoCA, we investigated group differences between CMN, CI, MI and CMI subjects using linear regression model given by: *MoCA* = *β*0+*β*_group_*group+*β*_sex_*sex+*β*_site_\**site*. For walk speed, the linear regression model given by: *Walk* = *β*0+*β*_group_*group+*β*_sex_\**sex*+*β*_YOE_\**YOE*+*β*_site_\**site*+*β*_BMI_*BMI, where *Walk* is the average raw walk speed for 2 trials of 4 meters walk speed. Sex, years of education (YOE), and BMI (body mass index) were accounted for in addition to *site* as categorical variable to account for the four HCP-A data acquisition locations. Bonferroni multiple comparison correction was done for the validation analysis.

To investigate microstructural group differences between CMN, CI, MI and CMI subjects, we used a linear regression model given by: *Pi*=*β*0+*β*_group_*group+*β*_sex_\**sex* +*β*_YOE_\**YOE*+*β*_site_\**site*+*β*_BMI_*BMI, where *Pi* is the mean ROI value of the parameter of interest (MAP metrics i.e., NG, PA, RTAP, RTOP, and RTPP and DTI metrics, i.e., FA, RD, AD, TR) of the *i*th ROI. The mean ROI MRI parameter, YOE and BMI were standardized using z-scores prior to regression. Results are presented as the beta coefficient of the *β*_group_ estimate which quantifies the group difference in the MRI variable, expressed in standard deviation units due to prior z-score normalization. False discovery rate (FDR) correction was done to correct for multiple comparisons [37] and the threshold for statistical significance was *p*_FDR_ < 0.05. Matlab was used for all computations.

## 3. Results

NIH-Fluid and walk endurances test scores [38] classified 582 subjects into 4 groups of CMN (n=113), CI (n=140), MI (n=195) and CMI (n=134). Table 1 presents participant demographics, test scores, and results of the group-wise ANOVA and Chi-square analysis. Age did not differ significantly between groups and was therefore not included as a covariate in the regression models. However, there was a significant difference in the distribution of males and females across the four groups, necessitating its inclusion as a covariate in the linear regression analysis of MRI metrics. Additionally, the CMN group had significantly less years of education compared to the CI and CMI groups. As expected, scores on the NIH-Fluid and Walk Endurance tests also showed significant group-wise differences.

### 3.1 Validation of group classification

To validate the study grouping, we analyzed MoCA and Walk Speed test scores from the HCP-A dataset. MoCA scores are commonly used for MCI screening and initial cognitive assessment [17]. Given that our classification was based on cognitive and motor performance, we used these measures as complementary and independent indicators alongside NIH-Fluid and Walk Endurance scores. A linear regression model was used to assess group-wise differences in MoCA scores, adjusting for sex and testing site as co-variates. As shown in Table 1, the CI and CMI groups had significantly lower MoCA scores compared to the CMN group. Similarly, a linear regression model was applied to the Walk Speed score, with sex, education, BMI, and testing site as covariates. The results indicated significantly lower walk speed performance in both the MI and CMI groups compared to the CMN group (Table 1).

### 3.2 Associations between cognitive-motor impairment and cerebral microstructure

This study utilized two dMRI models to characterize cerebral microstructure: the widely used DTI and the advanced MAP-MRI framework. Unlike biophysical model-based dMRI methods, MAP-MRI operates without prior assumptions, making it particularly effective for analyzing complex GM microstructures. Neither model (DTI or MAP) directly measures structural properties such as axon diameter [39] or neurite density [40] , requiring careful interpretation of derived metrics. In MAP-MRI, zero-displacement probabilities (RTAP, RTOP, RTPP) serve as proxies for cellular density and extracellular volume [41]. Additionally, PA captures diffusion directionality, while NG reflects microstructural heterogeneity by identifying regions with varying diffusivity profiles [35]. Table 2 summarizes the physical mechanisms and commonly accepted biological interpretations of these MAP-MRI parameters. Similarly, DTI provides a lower-order approximation of PA through FA, while the diffusivity parameters (AD, RD, TR) offer insights into different diffusion orientations and overall tissue integrity. By integrating these models, this study enhances the characterization of microstructural differences across cerebral regions.

To investigate brain regions associated with cognitive and motor impairment, we selected temporal-meta and motor-related ROIs, comprising 13 GM regions and 14 corresponding WM tracts (Figure 1). Axial and sagittal RTOP parameter maps for these regions, averaged across study participants, are shown in Figure 2 for the CMN, MI, CI, and CMI groups within restricted age ranges of 36–44, 45–55, 56–71, and 72–90 years. The total number of participants in each group is also reported. Although ANOVA results indicated no significant age differences between groups, specific age ranges were examined to qualitatively assess potential age-related effects on RTOP parameter maps. Visual inspection suggests that across all age ranges, participants in the CMI group generally exhibit lower regional RTOP values compared to other groups. Violin plots of z-scores of all MAP-MRI and DTI parameters across the four groups in representative ROIs of GM and WM are shown in Figure 3.

**Figure 2:**
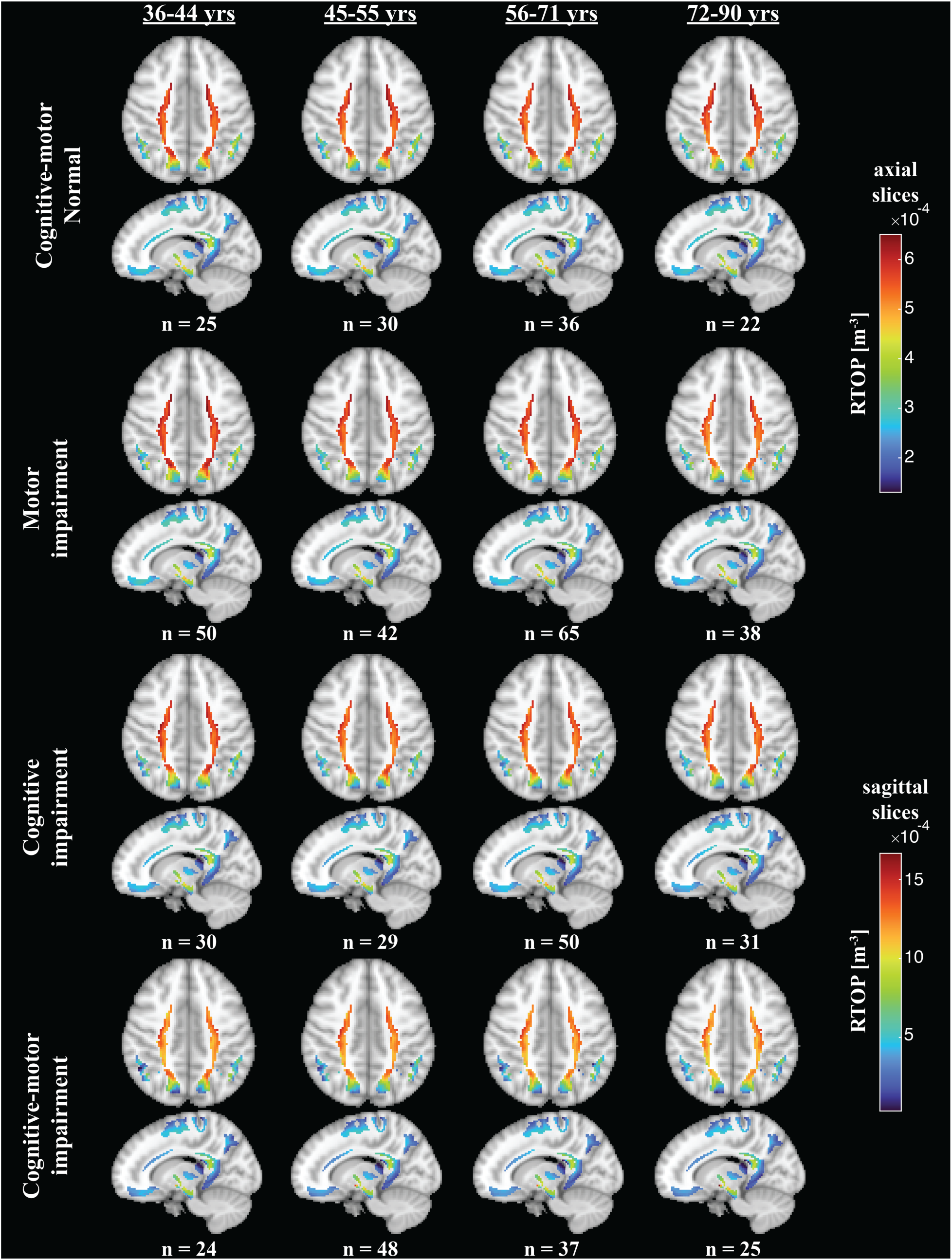
Representative MAP-MRI parametric maps. Examples of axial and sagittal RTOP parameter maps averaged across participants within the cognitive-motor normal (CMN), cognitive impairment (CI), motor impairment (MI), and cognitive-motor impairment (CMI) groups, at four age ranges of 36-44 years, 45-55years, 56-71 years and 72-90 years. The number of participants in each group is indicated (n). Visual inspection indicates that, overall, participants in the CMI group exhibit greater regional RTOP values, regardless of age.

**Figure 3:**
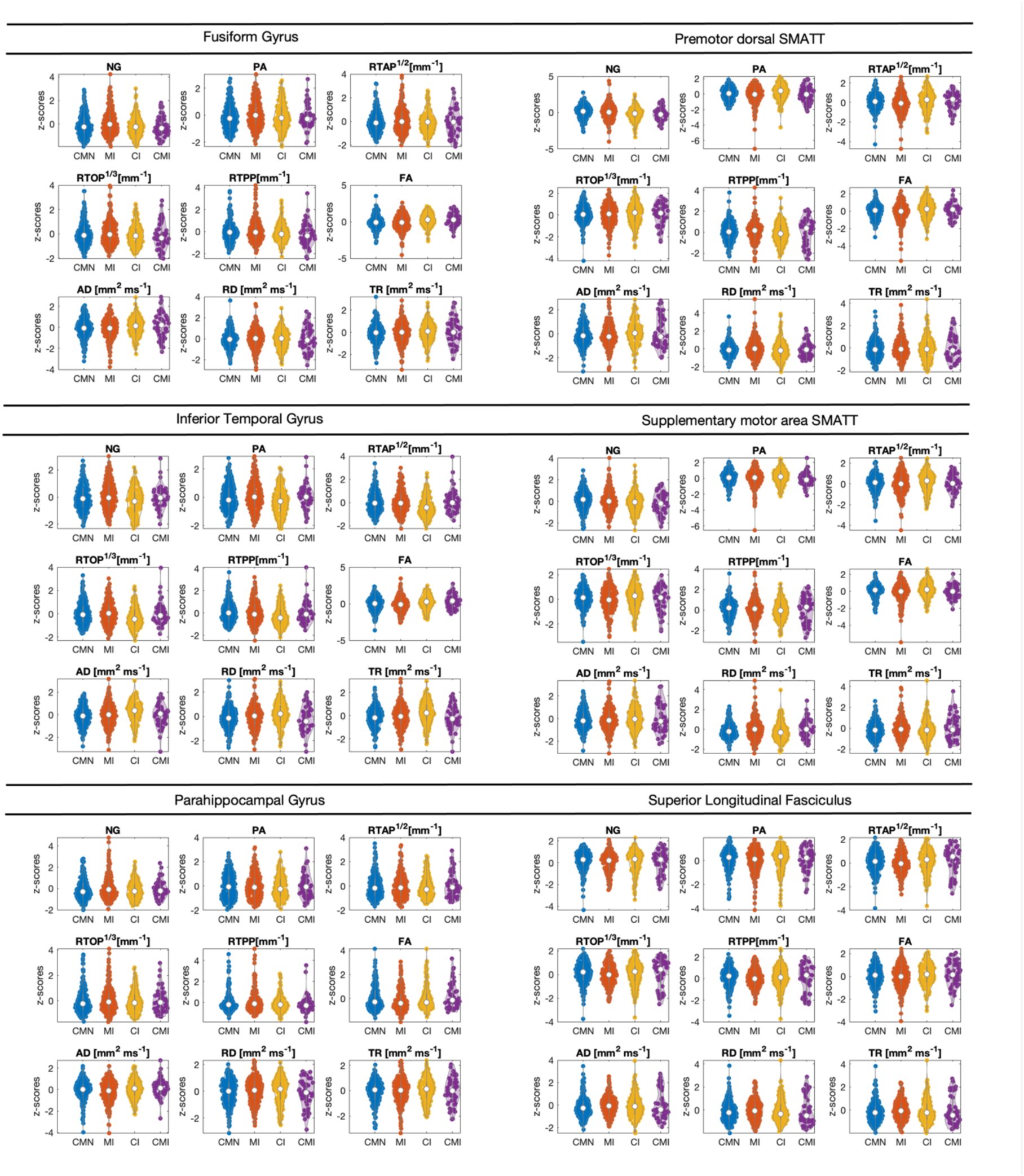
MRI parameters distribution across the different study groups. Violin plots showing z-scores of all MAP-MRI and DTI parameters across four groups in representative gray and white matter ROIs.

For each MRI metric and ROI, we show associations uncorrected and with FDR multiple comparison correction (*p*_FDR_ <0.05) to allow for exploratory analysis of our findings in Figures 4A and B, respectively. The results highlight the synergistic impact of dual cognitive-motor impairment on cerebral microstructure. Widespread microstructural differences between the CMN and CMI groups are evident after FDR correction, as indicated by the β coefficients in Figure 4B. In contrast, cognitive impairment alone (CI group) has a more limited effect on cerebral microstructure, with only a small number of affected ROIs remaining significant after multiple comparisons correction. Similarly, motor impairment alone (MI group) exhibits negligible microstructural differences when compared to the cognitively and motorically normative (CMN) group, none of which survives FDR correction.

**Figure 4:**
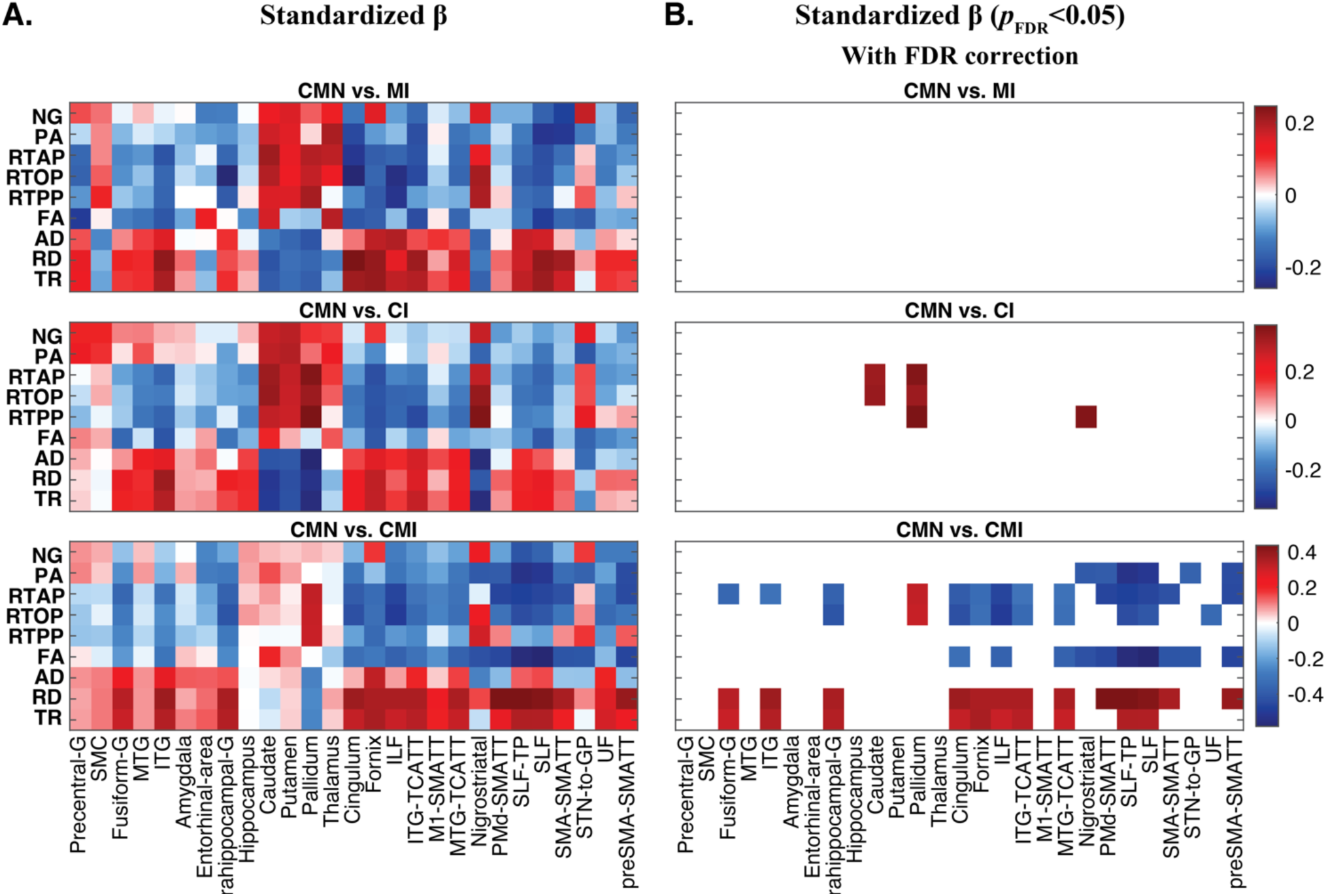
Region-specific associations of microstructural features and cognitive-motor impairment. Group differences of diffusion MRI metrics in 13 GM and 14 WM ROIs. (A) The β_group_ coefficients are shown as a matrix for diffusion MRI z-normalized features across all 27 ROIs. (B) Only p_FDR_ <0.05 β_group_ are shown. The CMI group exhibited significantly higher RTAP/RTOP/RTPP and lower RD/TR values compared with the CMN group in most WM regions and notably in GM regions of fusiform gyrus, inferior temporal gyrus and parahippocampal gyrus. CMN vs CI also showed significant difference in subcortical regions and nigrostriatal WM regions. CMN vs MI did not show any significant difference post multiple comparison correction. Abbreviations: sensorimotor cortex (SMC), inferior longitudinal fasciculus (ILF), inferior temporal gyrus (ITG), middle temporal gyrus (MTG), superior longitudinal fasciculus (SLF) and uncinate fasciculus (UF), inferior temporal gyrus transcallosal tract (ITG-TCATT), M1motor area tract - sensorimotor area tract template (M1-SMATT), middle temporal gyrus transcallosal tract (MTG-TCATT), nigrostriatal tract, dorsal premotor (PMd-SMATT), superior longitudinal fasciculus-tapetum (SLF-TP), supplementary motor area (SMA-SMATT), substantia nigra-globus pallidus (STN-GP), presupplementary motor area (preSMA-SMATT).

Specifically, we identified significant effect sizes in MAP-MRI-derived PA, RTAP, and RTOP, and in DTI-derived RD and TR, when comparing individuals with dual cognitive-motor impairment to the CMN group. These differences were observed in temporal-meta GM regions, such as the fusiform gyrus, ITG, and parahippocampal gyrus. Additionally, significant differences were found in WM relating to both temporal-meta and motor regions, including the cingulum, fornix, ILF, UF, ITG, nigrostriatal, and MTG-tracts, and several sensorimotor area tracts. FA showed similar pattern as zero-displacement probabilities in all WM regions (except UF) and in the GM region of ITG. Nearly all examined regions exhibited negative associations in PA, RTAP, RTOP, and FA, indicating significantly lower values in the CMI cohort compared to the CMN group. In contrast, TR and RD parameters displayed the opposite trend, with higher values in the CMI group. This pattern aligns with expectations, as diffusivity is inversely related to zero-displacement probabilities.

When comparing the cognitive impaired and the cognitive-motor normative groups, significant effect sizes were limited to the caudate, pallidum and the nigrostriatal tracts, and were only reflected in the MAP-MRI-derived zero-displacement probability parameters. No microstructural differences between the motor-impaired and the cognitive-motor normative groups survived FDR correction.

Although in most cases not surviving FDR correction, subcortical regions exhibited opposing microstructural trends compared with WM regions in all groups (Figure 4A). Specifically, higher NG, PA, and zero displacement probability values, and lower diffusivity values, in all the impairment cohorts compared to the CMN group.

## 4. Discussion

Through neuroimaging analysis of brain aging, we identified distinct neurobiological characteristics in adults experiencing dual impairment in cognition and motor function. Compared to individuals without cognitive or motor impairments, the dual impairment group exhibited extensive microstructural alterations indicative of reduced integrity, particularly in temporal meta-ROIs, motor-related GM regions, and associated WM tracts. While studies on dual impairment have primarily focused on macroscopic GM changes [42] or DTI-based WM assessments [7], the advancement of MAP-MRI now allows for GM microstructural investigation without model-specific assumptions. Our findings align with prior DTI studies on association tracts [7] and further advance spatial patterns of microstructural alterations in GM. Notably, only the dual impairment group showed widespread reductions in cellular density and increased extracellular volume across key GM and WM regions, highlighting potential mechanisms underlying dual impairment and its potential progression to dementia.

To assess cognitive, motor and dual impairments in the HCP-A cohort, we utilized NIH-Fluid and Walk Endurance measures. Fluid ability reflects learning and processing capacity, peaking in early adulthood and declining with age. Walk Endurance, linked to brain volume [43], cognitive function [44], and cardiovascular health [45], may enhance the sensitivity of gait-based classifications for MCI and dementia [46]. We classified participants into four groups based on impairment in these domains, using Walk speed and MoCA scores for validation. Regression analysis confirmed significant differences in cognition and motor function, supporting our classification approach.

Among the impairment groups, only the dual impairment group exhibited altered zero-displacement probabilities and diffusivity parameters in GM regions. These differences were observed in key temporal meta GM regions, including the parahippocampal gyrus, inferior temporal gyrus, and fusiform gyrus. Moreover, extensive differences between the dual impairment and control groups were identified across nearly all analyzed WM tracts. The dual impairment group exhibited lower PA and zero-displacement probabilities, along with higher diffusivity values, suggesting reduced fiber orientation coherence and potential demyelination. Put together, these findings suggest a higher degree of etiological heterogeneity within the dual impairment group. The cognitive impairment group exhibited differences from the CMN group in only a few regions, demonstrating higher zero-displacement probabilities—an inverse trend compared to most WM regions—specifically in the caudate, pallidum, and nigrostriatal tract. In addition, there were no significant microstructural differences between the motor impairment and the CMN group in any of the regions.

This study examines microstructural changes within tau PET-derived meta-ROIs, a set of regions that capture a broad dynamic range from normal aging to AD dementia, focusing specifically on their role in the prodromal stages of dual impairment [47]. Specifically, the entorhinal cortex, which relays cortical information from association cortices—either directly or through the perirhinal and parahippocampal cortices—to the hippocampus and other limbic structures, plays a crucial role in memory and cognitive processing [48]. The microstructural changes observed in the inferior temporal gyrus, parahippocampus, and limbic-associated tracts (cingulum, ILF, SLF) in the dual impairment group may reflect disruptions in these pathways, potentially contributing to deficits in executive function, cognitive flexibility, attention, episodic memory formation, and processing speed, regardless of age. The premotor cortex and supplementary motor area play a critical role in selecting, planning, and coordinating voluntary movements, including gait, and therefore the observed microstructural changes in motor tracts of the PMd, SMA, and preSMA may be indicative of reduced endurance and speed. Given the established link between slower walking speed and cognitive impairment—along with its predictive value for long-term cognitive decline [49] these findings suggest that the dual impairment group experiences a compound effect of cognitive and motor decline.

Our findings reveal a distinct microstructural pattern in the examined subcortical regions, characterized by higher NG, PA, and zero-displacement probability values, along with lower diffusivity values, across all impairment cohorts compared to the CMN group. While most of these associations did not remain significant after FDR correction, the spatial specificity of this trend warrants further exploration. The complex architecture of these subcortical regions, particularly the presence of crossing fibers [50–52], provides context for interpreting these microstructural alterations. The observed increases in NG, PA, and zero-displacement probabilities suggest reduced neurite orientation dispersion, which we hypothesize is driven by a decline in crossing fibers and diminished regional connectivity.

Supporting our results, a recent longitudinal DTI study showed that a dual cognitive-motor decline group undergoes faster FA reduction in commissural and association tracts, yet no significant differences were observed cross-sectionally [7]. However, the study did not examine GM microstructural characteristics or WM tracts specifically associated with temporal meta-ROIs. The findings we present here align with previous research demonstrating microstructural alterations in the inferior temporal gyrus, where amnestic MCI and AD cases exhibit a significant reduction in synaptic density (36% fewer synapses) compared to cognitively intact individuals [53]. Synaptic loss in this region has been strongly correlated with Mini-Mental State Examination scores and category verbal fluency performance [54]. The observed inverse relationship between the dual cognitive-motor impairment and zero-displacement probabilities is consistent with prior MAP-MRI studies [13,16] and reflects established patterns of GM microstructural changes across the adult lifespan [55]. These results suggest that CMI individuals may experience decreased cellular density and increased extracellular volume in GM. Additionally, neuroinflammation—a hallmark of neurodegenerative pathology—triggers the activation of microglia and astrocytes, leading to morphological changes such as astrocytic hypertrophy, characterized by cell body expansion and elongated processes [56]. These glial alterations may contribute to the observed reductions in MAP-MRI-derived zero-displacement probabilities in the CMI group.

Despite its strengths, this study has several limitations. While the cohort is well-characterized, it was not specifically designed to track cognitive and motor changes over time (i.e., longitudinally), limiting its ability to precisely detect dual decline and dementia progression. The absence of FLAIR imaging and comorbidity data prevents a comprehensive assessment of cardiovascular contributions to gait slowing, cognitive decline, and executive dysfunction [57]. However, the large sample size from a general voluntary population remains a notable strength. Previous research has primarily focused on volumetric GM changes due to constraints in existing biophysical models. To the best of our knowledge, this study is the first to apply the MAP-MRI model, a diffusion MRI framework independent of biophysical assumptions, to assess GM microstructural changes associated with cognitive-motor impairment in the HCP-Aging cohort. The HCP-A maximal diffusion encoding strength (b=3000 s/mm^2^) allows using a MAP-MRI expansion of 22 coefficients (order 4), while extending to order 6 by incorporating higher b-values (up to 6000 s/mm²) [58] is considered optimal for clinical applications [35]. While the linear diffusion encoding used here cannot distinguish between microstructural size and orientation, alternative methods such as planar [59] or spherical [60] encoding may address this limitation. Additionally, integrating relaxation and diffusion encoding is recommended to enhance sensitivity and specificity [61,62].

In summary, dual cognitive-motor impairment may serve as an aging phenotype, characterized by diminished cellular density in temporal GM meta ROIs, lower fiber orientation coherence and potential demyelination in associated WM tracts, and by attenuated regional connectivity in subcortical regions. Continuous monitoring of both cognitive and motor function is essential for identifying older adults exhibiting dual impairment, as this may indicate an increased risk of neurodegenerative processes leading to dementia. Early identification of these individuals allows for timely diagnostic assessments to determine underlying neuropathological changes. Our findings highlight the need for longitudinal studies on dual impairment utilizing multi-shell diffusion MRI, which enables the assessment of GM microstructure. Such investigations are essential for tracking within-individual changes over time and gaining a deeper understanding of brain aging trajectories. The development of new therapeutic approaches presents an opportunity to implement targeted interventions for individuals experiencing dual cognitive-motor impairment, potentially reducing the progression of neurodegeneration and postponing the onset of dementia.

## Consent Statement

Consent was obtained according to the protocols approved by the Institutional Review Board (IRB).

